# Sensitivity to Cuticular Hydrocarbons Across the Odorant Receptor Family in the Indian Jumping Ant

**DOI:** 10.1101/2025.07.17.665333

**Authors:** Røskva Tórhalsdóttir, Benjamin I. Morris, Aiden Masters, Bogdan Sieriebriennikov, Amatullah Tawawalla, Lydia F. Naughton, Deanna Cannizzaro, Jenna Longo, Kendall Ham, Bri Pomonis, Alex Lyford, Jocelyn G. Millar, Gregory M. Pask

## Abstract

Eusocial insects rely on the olfactory detection of cuticular hydrocarbons (CHCs) to mediate fundamental eusocial behaviors, such as nestmate recognition, and reproductive division of labor. In the ponerine ant *Harpegnathos saltator*, highly expanded odorant receptor (OR) families detect CHCs and mediate these eusocial behaviors at the molecular level. Previous studies have characterized *H. saltator OR* (*HsOr*) genes within the 9-exon and other large subfamilies, but it remains unclear how other *HsOr* subfamilies may contribute to CHC detection. Using heterologous expression in *Drosophila melanogaster* olfactory neurons, we characterized HsOr sensitivity more broadly across the gene family, outside the 9-exon subfamily, to a panel of hydrocarbons (HC). Twenty-three HsOrs across sixteen subfamilies were screened, and several were found to be broadly tuned and weakly responsive to the HCs tested, except for HsOr152 which showed narrow tuning to a single HC found on the *H. saltator* cuticle. Lastly, we compiled and analyzed the HC responses from the 70 HsOrs from this and previous studies. This analysis suggests a combinatorial coding model of CHC detection, where several receptors across different subfamilies can contribute to the detection and discrimination of different CHCs. Our characterization of HsOrs provides functional insights into the molecular mechanisms of chemical communication among eusocial insects.

## INTRODUCTION

In eusocial insects, chemical communication among individuals in a colony influences key social behaviors such as nestmate recognition and reproductive division of labor (1). An important class of chemicals that mediates these social interactions is cuticular hydrocarbons (CHCs), mixtures of long-chain alkanes and alkenes which coat the hydrophobic epicuticle of terrestrial insects and other arthropods (2). CHCs primarily serve as protective barriers against desiccation and abrasion, but eusocial Hymenoptera and other social insect taxa have co-opted and diversified CHC profiles to signal task allocation, discriminate nestmates from non-nestmates, and suppress ovary development in sterile workers (3–6).

In the Indian jumping ant, *Harpegnathos saltator*, the roles and detection of CHCs have been previously examined at several levels. *Harpegnathos saltator* is a primitively eusocial ant species in the subfamily Ponerinae that has relatively small colony sizes (65 ± 40 workers) and a unique social structure defined by reproductive plasticity (7). When a queen’s fecundity decreases, workers in the colony can transition to a reproductive caste, termed a gamergate, and take over egg-laying duties. During the worker-to-gamergate transition, their CHC profiles shift to longer chain hydrocarbons which resemble those of reproductive queens (8). Notably, 13,23-dimethylheptatriacontane (13,23-DiMeC37) is the largest CHC by mass found in the species and is only present in reproductive queens and gamergates (8). It is hypothesized that 13,23-DiMeC37 and a subset of other CHCs constitute a fertility signal which serves to suppress ovarian activity and gamergate transition in workers.

CHCs in *H. saltator* and other ants are detected by basiconic sensilla on the antennae. These sensilla are only present in female castes, house over >50 olfactory receptor neurons, and are increasingly abundant in the distal segments of the antennae (9,10). In *H. saltator* and ants in the *Camponotus* genus, electrophysiological analysis of basiconic sensilla revealed that they are responsive to a wide range of CHCs (11–13). Additionally, *H. saltator* gamergates showed decreased sensitivity to several long-chain CHCs when compared to workers, suggesting that an olfactory shift accompanies their increase in reproductive activity (11).

To understand the molecular basis of CHC detection, attention has turned to the odorant receptors (ORs) in eusocial insects. Genomic analyses in ants have revealed greatly expanded *OR* families in ants compared to other insects, with >300 *ORs* identified across several species (14–17). Notably, the 9-exon subfamily of ant *ORs*, which represents roughly a third or more of the entire family, is highly enriched in antennal transcriptomes, and is under significant positive selection (16–19). Recent studies using heterologous expression in *Drosophila melanogaster* ORNs have shown that *H. saltator* ORs (HsOrs) are sensitive to HCs, both individually and as mixtures in cuticular extracts. The 9-exon subfamily HsOrs showed variable tuning to HCs, with some broadly responsive to several HCs and others sensitive to only a single or a few structurally similar HCs (20). Interestingly, HC sensitivity was also observed in several other HsOr clades, suggesting that despite the prominence of 9-exon HsOr genes, HC detection may rely on combinatorial coding that extends beyond this expansive subfamily (21).

In this study, our objective was to explore HsOr tuning to HCs broadly across the 381 HsOr family. Through expression in *Drosophila*, we functionally screened 23 HsOrs against a panel of HCs, many of which are found in *H. saltator* cuticular extracts. We then extended our analyses to include all existing HsOr functional data to observe trends in coding capacity relative to HsOr subfamily and HC chain length. With a total of 70 HsOrs screened across this HC panel, this work provides insights into the combinatorial coding that can facilitate chemical communication across social insects.

## RESULTS

HsOr functional responses were assessed against a panel of both commercially available alkanes and synthesized methyl-branched and unsaturated HCs as previously described (20,21). A total of 23 HsOr genes were either cloned from antennal cDNA or synthesized based on sequences from antennal transcriptomes (Figure 1A) (16). These HsOr genes were chosen to cover a broad range of HsOr genes from the 20 subfamilies outside of the expansive 9-exon subfamily that was previously characterized (20). During the selection process, priority was given to HsOr genes with higher transcript abundance in worker antennae relative to males (Figure 1A, B). We also included two genes from very small subfamilies – HsOr62 from subfamily M and HsOr219 from subfamily C, even though their expression was not female-biased. Finally, we included HsOr16 to compare it to the very closely related HsOr13, even though HsOr16 also did not show female-biased expression.

**Figure 1.**
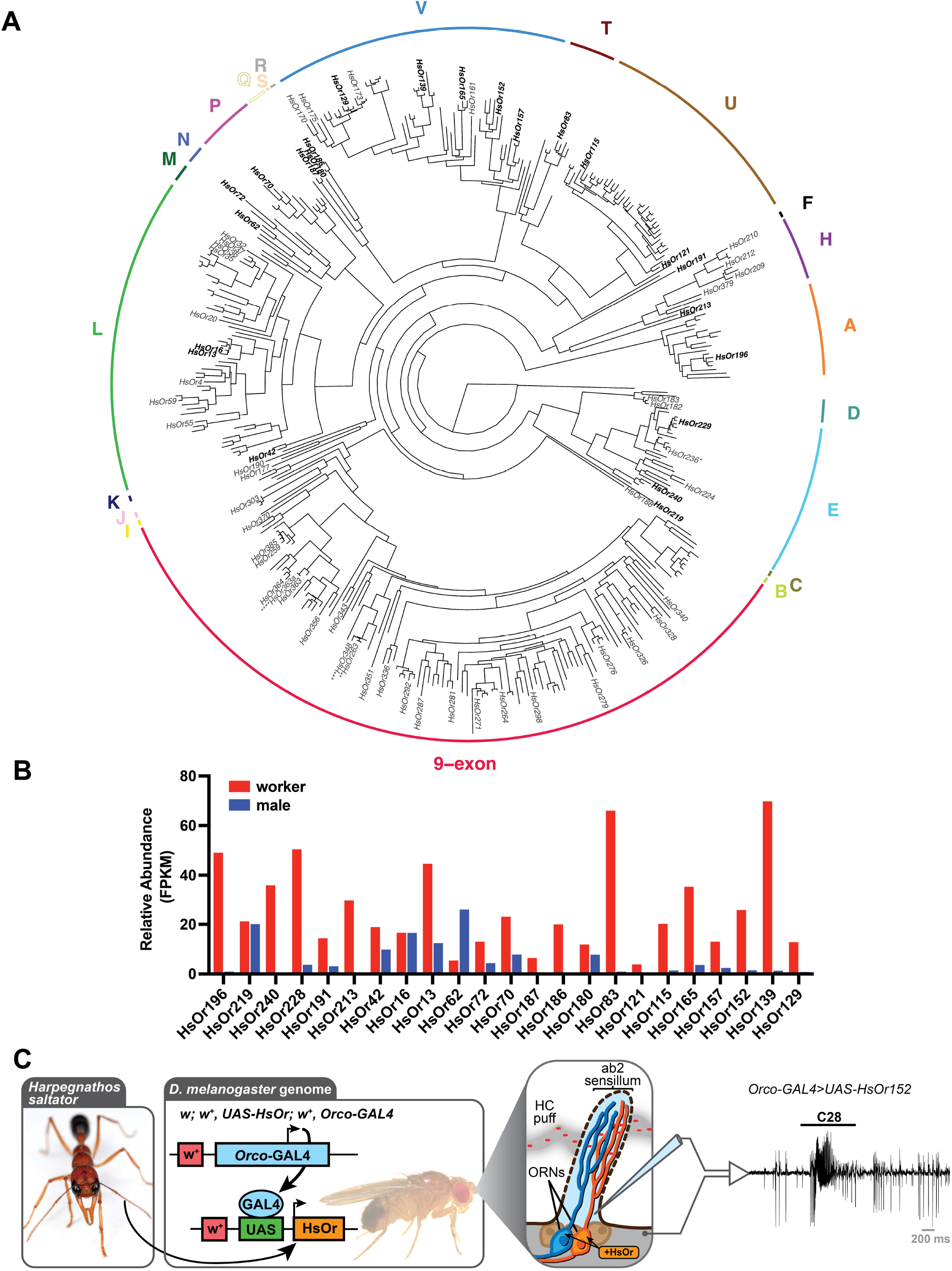
Functional characterization of HsOrs from across the gene family. (A) Phylogenetic tree of the full HsOr family with labeled gene names from previously characterized HsOrs. HsOrs decoded in this paper are shown in bold. Updated annotations of HsOrs are indicated by asterisks with more details in the Methods section. *HsOr236 sequence in Slone et al. 2017 was synthesized based on old gene annotations and represents a hybrid between current HsOr236 and HsOr238 (21). **HsOr263 cloned sequence from Pask et al. 2017 likely represents a truncated version of HsOr348 (20). ***HsOr348 sequence in Pask et al. 2017 was synthesized based on old gene annotations and represents a hybrid between current HsOr347 and HsOr348 (20). ****HsOr363a sequence in Pask et al. 2017 is a splice-isoform of HsOr363 (20). (B) Expression values in fragments per kilobase per million reads (FPKM) from transcriptomics data from males and workers (16). (C) Schematic for HsOr cloning, *in vivo* heterologous expression in *D. melanogaster* using the GAL4-UAS binary expression system, and electrophysiological recordings from an ab2 sensillum of a *w*; *w*^*+*^, *UAS-HsOr152; w*^*+*^, *Orco-GAL4* fly in response to an air puff through a heated cartridge dosed with 20 nmol of octacosane (C28). Inset *H. saltator* image is provided courtesy of Alex Wild (alexanderwild.com).

Heterologous expression of HsOr genes in olfactory receptor neurons (ORNs) of *D. melanogaster* was achieved by generating a *UAS-HsOr* transgenic fly and crossing it with a robust *Orco-GAL4* driver line (Bloomington Drosophila Stock Center #23292). As done previously, HsOr function was assessed using single-sensillum recordings (SSR) from the ab2A neuron and a screening panel of 39 HCs, of which 17 have been found in *H. saltator* cuticular extracts (Figure 1C) (8,20,21).

From this dataset of 897 distinct HC-HsOr pairs, we found that eleven of the 23 HsOrs responded to at least one HC above 30 Δspikes/s, a threshold set six times higher than the native ab2A spontaneous firing rate (Figures 2 and 3) (22). All of these >30 Δspikes/s responses were elicited by HCs with a chain length of C21 or longer, indicating a slightly lower detection range relative to the previously characterized HsOrs where no responses were found below C27 (20,21). Six HsOrs (HsOr240, HsOr191, HsOr213, HsOr70, HsOr157, HsOr139) demonstrated broad tuning with >30 Δspikes/s responses to seven or more HCs on the panel. Twelve HsOrs did not show a >30 Δspikes/s to any HC on the panel, with four (HsOr219, HsOr187, HsOr165, HsOr129) having generally weak negative responses across the panel where the HCs inhibited the firing rate of HsOr-expressing ab2A neuron.

**Figure 2.**
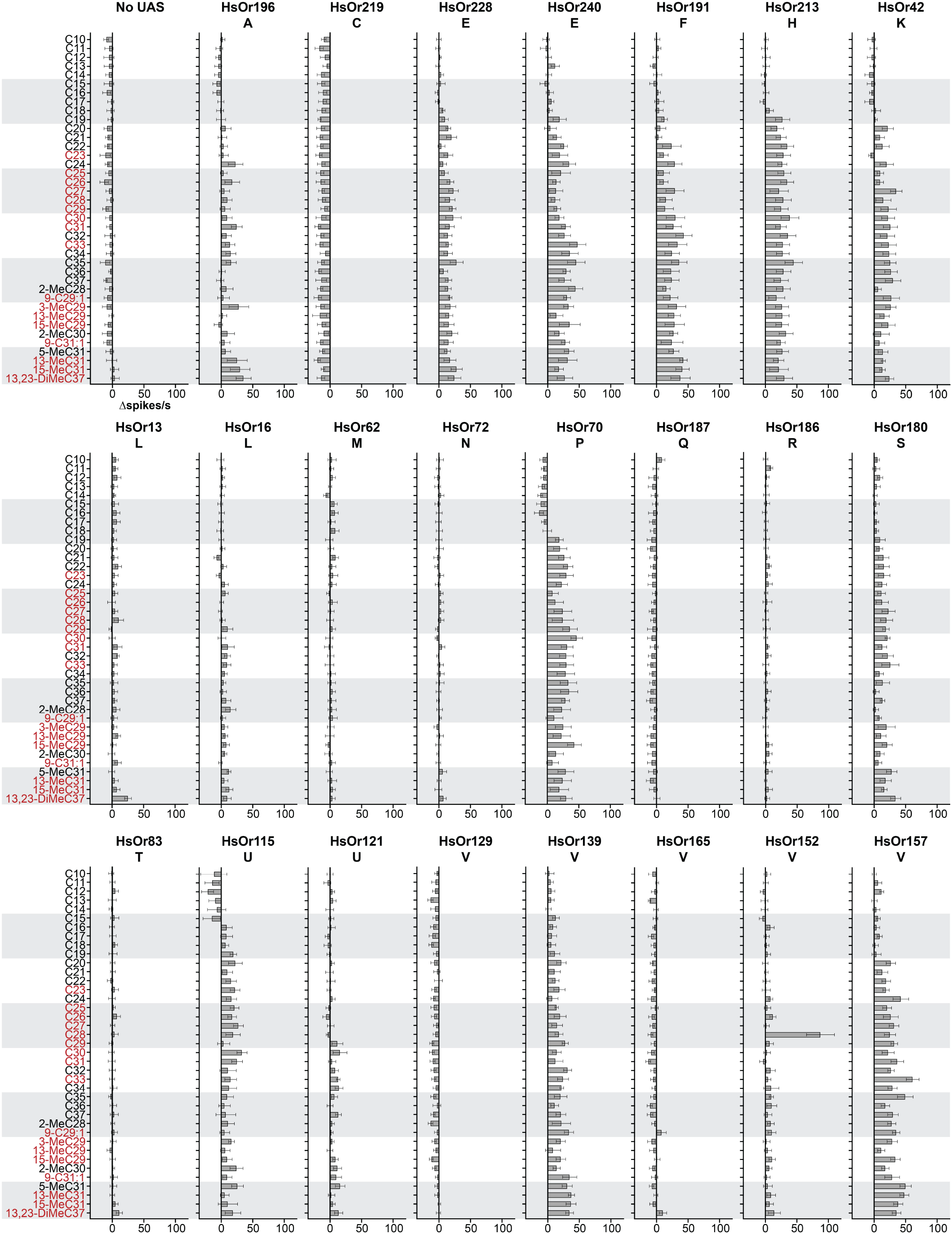
HC detection across the several subfamilies of HsOrs. Mean responses (in Δspikes/s) of each HsOr to a panel of 39 HCs (each at 20 nmol) from the ab2A neuron of *w*; *w*^*+*^, *UAS-HsOr; w*^*+*^, *Orco-GAL4* flies. Values represent mean□±□SEM (*n*□=□6). Respective subfamily is denoted below HsOr name, and HC names in red are present in *H. saltator* cuticular extracts (8).

**Figure 3.**
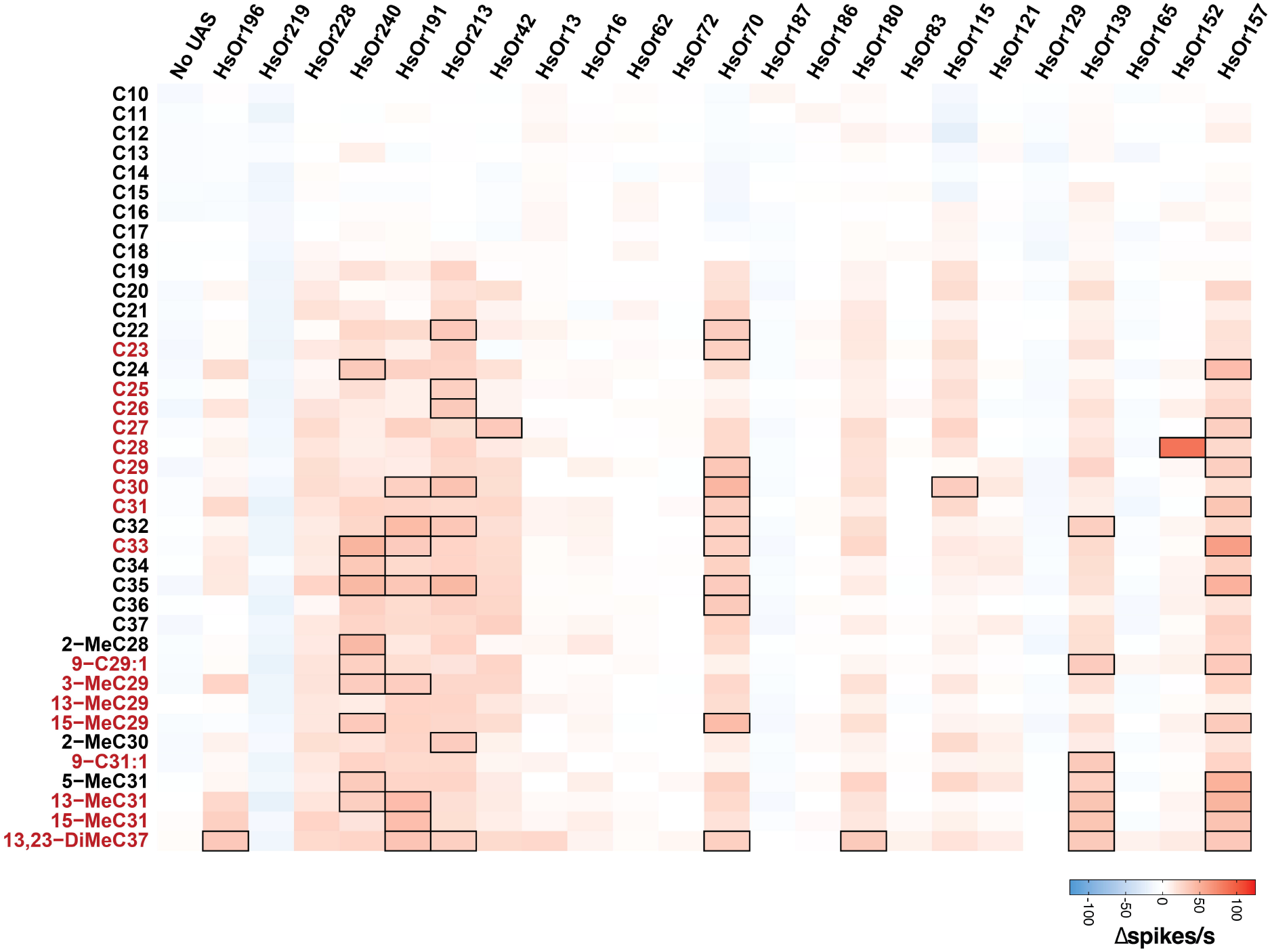
Heat map of mean HsOr responses in Δspikes/s to the HC panel. A heat map representation of data in Figure 2. HC names in red are present in *H. saltator* cuticular extracts (8). Black borders indicate mean responses above the 30 Δspikes/s threshold. HC names in red are present in *H. saltator* cuticular extracts (8).

One receptor, HsOr152 of the V subfamily, was found to be narrowly tuned to C28, with a mean response of 87.5 Δspikes/s (Figures 1C, 2, and 3). Notably, two other V subfamily receptors, HsOr129 and HsOr165, displayed largely inhibitory responses to many HCs on the panel, with several pairs that resulted in -10 Δspikes/s or greater inhibition of the ab2A spontaneous firing rate. With the exception of HsOr152, most receptors showed overall weaker and broader tuning to HCs relative to the previously decoded 9-exon HsOrs (20,21).

To further explore the HC sensitivity, we performed dose-response assays for the most efficacious HsOr-ligand pairs for the 11 HsOrs with a >30 Δspikes/s response (Figure 4). Dose response curves showed that HC detection occurred in a concentration-dependent manner (Figure 4). Interestingly, several HsOrs had notably larger (HsOr196, HsOr191, HsOr157) or smaller (HsOr240, HsOr152) responses to 20 nmol of HC when compared to the same dose in the screening panel, suggesting that the repeated HC stimulation during the screen may positively or negatively affect receptor response dynamics. Additionally, no HsOrs showed any responses above 10 Δspikes/s at or below 2 nmol of HC.

**Figure 4.**
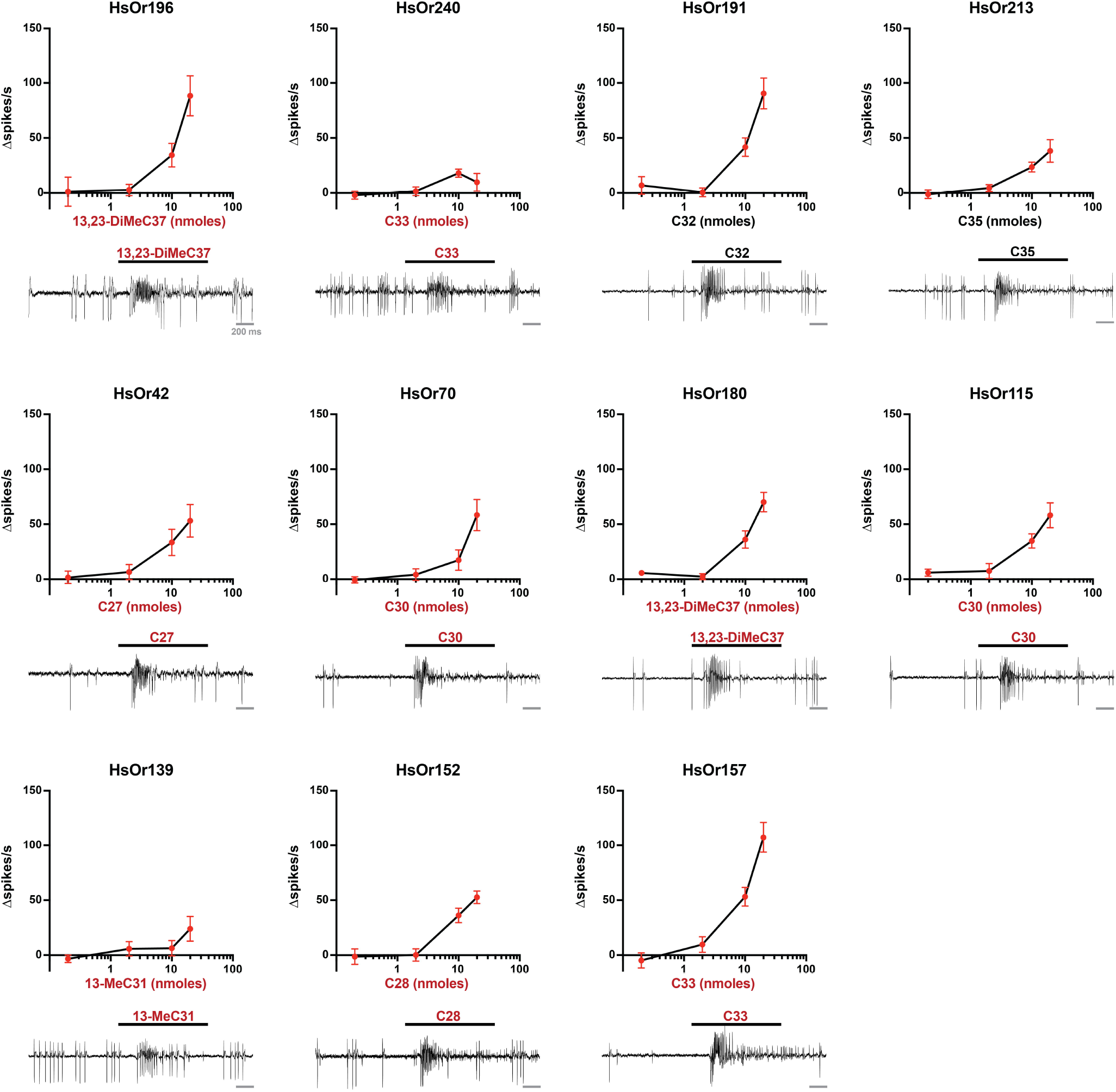
Concentration-dependent responses of HsOrs. (top) Dose-dependent curves (top) and a representative excitatory response at 20 nmol (bottom) to the most efficacious HC ligand for each HsOr that displayed responses above 30 Δspikes/s in the initial screen (mean□±□SEM, *n*□=□6). HC names in red are present in *H. saltator* cuticular extracts (8).

## DISCUSSION

The 23 HsOrs characterized in this study demonstrate that HC detection is not limited to the greatly expanded 9-exon subfamily as found previously (21). HC response profiles varied greatly across and within subfamilies, but overall showed broader tuning and relatively weaker responses for most HsOrs in this study compared to the 9-exon study (excluding HsOr152). Although some HC-evoked responses were below our 30 spikes/s threshold, it is still possible that they may play a role in the combinatorial coding of HCs in the *H*. saltator antennae, where a single HC can elicit responses from multiple chemosensory receptors. The characterized HsOrs displayed a preferential tuning to longer chain HCs above C21, which can be attributed to their ecological relevance as several of these have been found in cuticular extracts from *H. saltator* (8). The panel used in this study did not contain all the CHCs found in *H. saltator* cuticular extracts. Thus, it provides a gateway into understanding HC detection and may not contain all the true ligands of the characterized receptors.

The narrow ligand specificity of some of these HsOrs was evidenced by the strong response of HsOr152 to C28 but weak response to C27, C29, and 2-MeC28. Similarly, a one-carbon difference in HC length between C29 and C30 resulted in either a lack of response (3.8 Δspikes/s) or a substantial excitatory response (33.0 Δspikes/s), respectively, in HsOr115. This discriminatory power indicates that these HsOrs can distinguish between small structural differences between HCs, suggesting that a deeper exploration of additional methyl-branched and unsaturated HCs, preferably those found in cuticular extracts, may yield additional ligand-receptor pairs.

Furthermore, four HsOrs (HsOr219, HsOr187, HsOr165, HsOr129) showed broad inhibitory responses to most HCs on the screening panel. This inhibition may play an important role in olfactory system modulation at the receptor level of the periphery, as previously observed in *Drosophila* (23). Here, the presence of salient HCs could actively reduce the ability to detect non-pheromone ligands through receptor-mediated inhibition. This mechanism could favor the detection of social cues through other receptors while turning down the “gain” of environmental stimuli.

Notably, the V subfamily showed a high degree of diversification and specialization with receptors being a combination of narrowly tuned (HsOr152), inhibited (HsOr129 and HsOr165) and broadly excited (HsOr157 and HsOr139) by HCs. Specifically, many of the >30 spikes/s responses of HsOr157 were evoked by odd-numbered carbon chain lengths, reflecting a similar pattern found in *H. saltator* CHC extracts, which are strongly biased towards odd-numbered chains. This reflects their biosynthesis, which consists of the sequential attachment of two-carbon acyl-CoA units, and culminating in a decarboxylation to give the odd-numbered chains (2,8). The wide functional variation within the V subfamily HsOrs aligns with the reported rapid expansion and diversification observed at the genetic level of the ant ORs, allowing them to display strikingly different ligand affinity and efficacy (18).

Many of the dose-response curves (Figure 4) showed less potent responses to the tested HCs when compared to the previously tested 9-exon HsOrs. Specifically, 2 nmol of the most efficacious ligand did not elicit a notable response. Additionally, some responses from the dose response data are higher or lower than those from the full screening panel. This could be due to inherent variability in heterologous *in vivo* expression, but it is also possible that sequential stimulation with the full HC panel could cause HsOr-expressing ORNs to be sensitized or desensitized, as has been observed in other insects, after exposure to structurally related HC ligands (24–26).

Twelve HsOrs did not respond to any of the HCs on the screening panel (<30 spikes/s). Possible explanations for these results include that these HsOrs may detect non-HC ligands, be narrowly tuned to a HC not in our screening panel, or be difficult to express in our heterologous *Drosophila* expression system. An expanded screening panel or expression in cell culture-based heterologous systems could address some of these possibilities.

With the addition of these 23 HsOrs, a total of 70 HsOrs have now been tested against a largely overlapping HC panel. To make these 70 HsOr response profiles accessible, we have developed a web application, the HsOr Response Database (https://pask-lab.shinyapps.io/HsOr_Response_Database/) to serve as a data repository. Overall, the functional data from across the broader HsOr family shows similar trends of generally weaker responses and broader tuning as compared to those in the 9-exon HsOrs (20,21). Combining all functional data by HsOr subfamily, we saw the greatest excitatory responses to straight-chain, long HCs between C28-37 (Figure 5). This mirrors the CHC profiles previously identified in *H. saltator* cuticular extracts, where 66 of 71 CHCs have a 28-37 carbon chain length (8). Given the CHC diversity in this range, it is possible that the plethora of methylated, dimethylated, and unsaturated HCs not in our screening panel may elicit greater responses from our characterized HsOrs. It is also important to note that the seemingly reduced responses to the methyl-branched alkanes from HsOrs outside the 9-exon subfamily, specifically the E, H, L, and V subfamilies, can be a result of 22 out of 70 HsOrs only being tested against straight chain alkanes (as shown in Figure 6A). Additionally, the two unsaturated HCs in the panel, Z9-C29:1 and Z9-C31:1, though present in *H. saltator* cuticular extracts, have yet to elicit notable responses from the screened 70 HsOrs. Inhibitory responses were present in a few representative subfamilies, with more widespread inhibition observed for short-chain HCs (C10-C17). It is also possible that the true ligands for these receptors have yet to be identified and may come from non-HC general odorant chemical classes, similar to previously studied HsOrs (16, 21).

**Figure 5.**
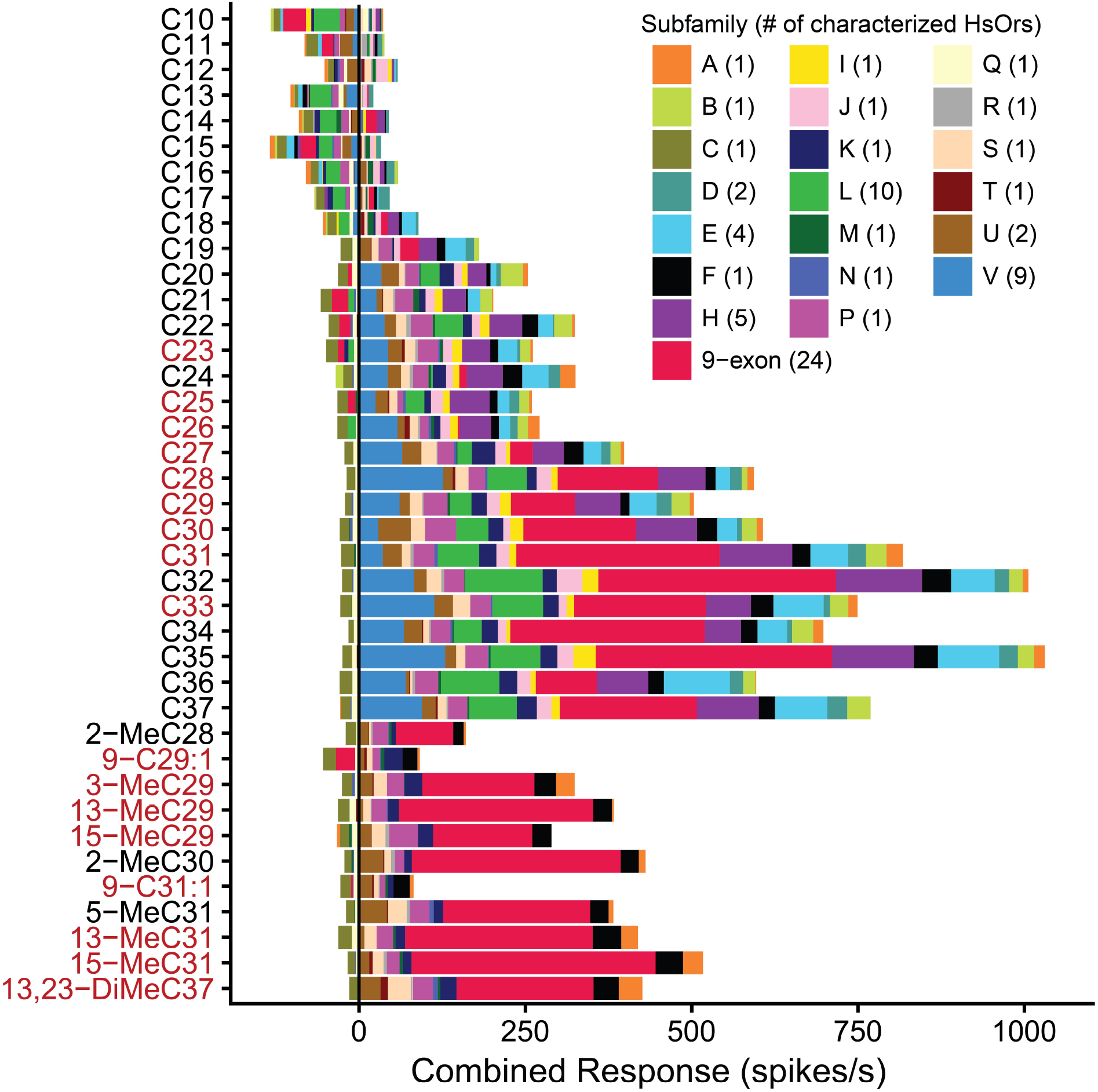
Combined HC response across all 70 decoded HsOrs. Each color represents different HsOr subfamilies and the number of decoded HsOrs in each subfamily is noted in the legend. HC names in red are present in *H. saltator* cuticular extracts (8).

**Figure 6.**
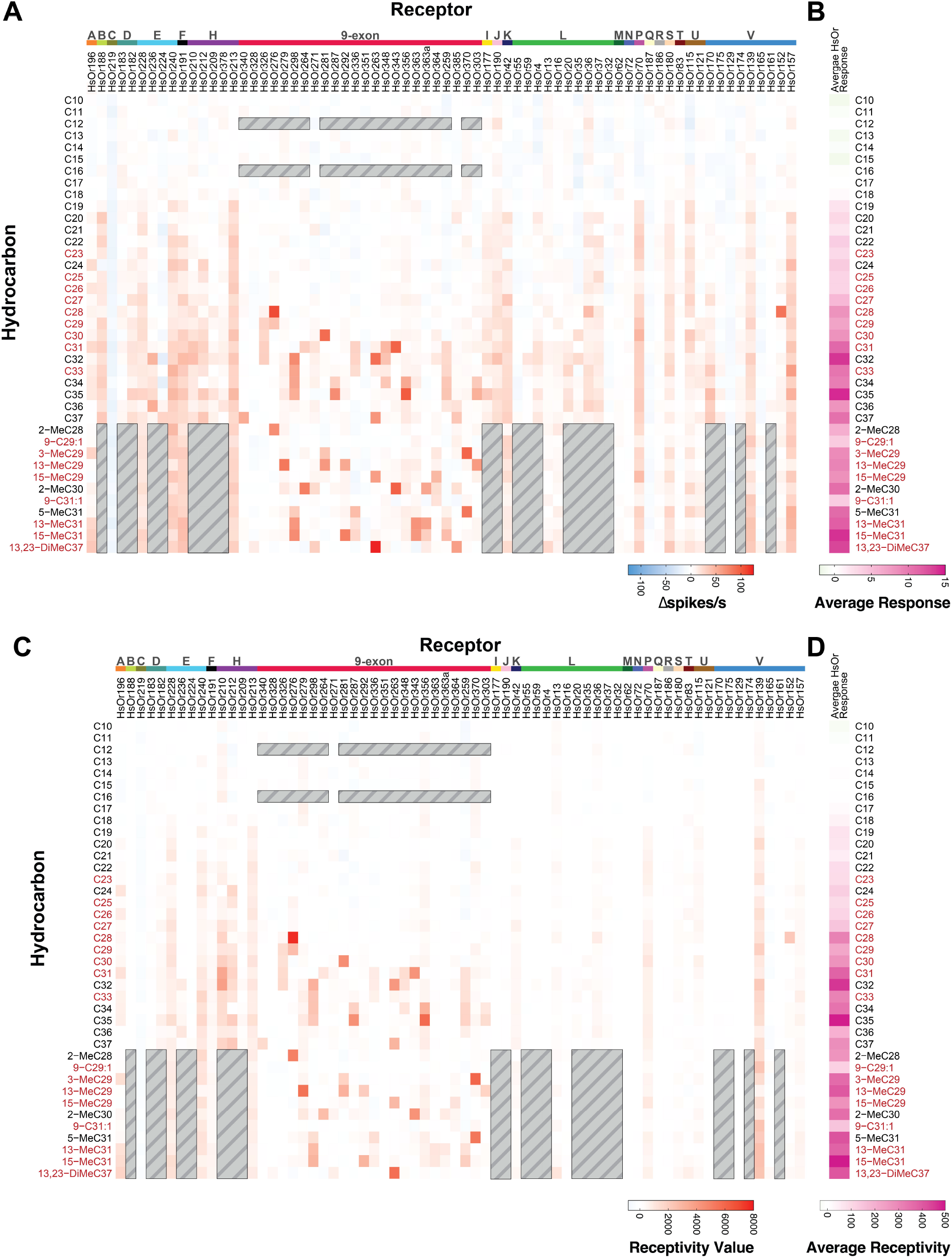
Mean and expression-weighted HsOr responses by HC. (A) Heat map of mean HsOr responses in Δspikes/s to the full panel of 39 HCs, each at 20 nmol. (B) The average response to each HC (summed Δspikes/s divided by the number of tested HsOrs). (C) Similar to panel A, but each response value is weighted by multiplying it by the corresponding HsOr expression value from worker antennae (in FPKM, from Zhou et al 2012 (16)) to produce a receptivity value (arbitrary units). HsOr 385 and HsOr378 were not included as no expression data was available. (D) The average receptivity to each HC (summed receptivity value to a HC divided by the number of tested HsOrs). For all panels, gray-lined boxes indicate HsOr-HC pairs that were not previously tested and HC names in red are present in *H. saltator* cuticular extracts (8).

We generated a heat map of all the 70 HsOr responses to the HC panel and then calculated the average receptor response for each HC (Figures 6A and B). The HCs with the six largest average responses were, in order, C35, 15-MeC31, C32, 13,23-DiMeC37, 13-MeC31, and 5-MeC31. We then generated a “receptivity” value for each HsOr-HC response by multiplying the spikes/s by each receptor’s previously reported expression value from a worker antennal transcriptome, as has been previously done with mosquito response data (Figure 6C) (16,27). When weighting these responses and averaging the receptivity for each HC, the top six receptivity values were 15-MeC31, C35, C32, 5-MeC31, 13,23 DiMeC37, and 13-MeC29. Previous electrophysiological recordings from basiconic sensilla of *H. saltator* workers showed responses to both straight-chain and methyl-branched HCs, but single-unit resolution is not possible given the >50 neurons housed in the basiconic sensilla (9,11). In the combined neuronal responses of basiconic sensilla, C10, C11, C23, C25, C28, and C32 elicited the six largest responses. While C28 and C32 elicited some large responses in our HsOr functional data, the other hydrocarbons did not activate any of the 70 characterized HsOrs, suggesting that other HsOrs in the odorant receptor family are responsive to these shorter chain HCs.

Overall, these functional data provide further evidence for the combinatorial coding involved in CHC detection by eusocial insects, where HsOrs can be either broadly or narrowly tuned to CHC ligands, and an individual CHC can bind to several HsOrs. We suggest that *H*. *saltator* can serve as a eusocial insect model for understanding the molecular basis of olfactory communication and its role in shaping social behaviors in a colony environment. While recent and further studies, such as the knockout of *Orco* in two ant species, are integral for a better understanding of the neurobiological mechanisms and behavioral outputs related to ant olfactory communication, functional characterization of these HsOrs can provide further insights into the HsOr-CHC interactions (28,29). Recent advances in cryo-EM 3D structures of insect ORs and protein modeling platforms can explore the observed specificity and/or flexibility of CHC-binding pockets with ant ORs (30–33).

## METHODS

### HsOr Gene Annotations and Phylogenetic Analysis

*OR* genes in *H. saltator* were annotated and assigned names by Zhou et al. (16). This naming convention was used by the previous papers that functionally characterized the ant *ORs* (20,21). However, this set of gene annotations was created for an older version of the genome assembly, which has since been superseded (34). Annotating *OR* genes in the new assembly and matching their names to the ones published by Zhou et al. took several rounds of manual curation (34,35). Yet, we still noticed several discrepancies between the old annotations, the new annotations, and the gene sequences used in the functional studies. We sought to correct them here. For example, the most recent version of gene annotations labeled as HSAL70 misses a gene labeled as *HsOr303*. We identified through BLAST search that *HsOr303* is the gene currently labeled as *LOC105187615* and renamed it to *HsOr303*. As another example, the new assembly is more contiguous, and the locus that contains *HsOr256, HsOr257, HsOr258, HsOr259, HsOr354, HsOr353*, and *HsOr352* contains two more *Or* paralogs that were absent from the old assembly and, as a consequence, from the old gene annotations. The sequence of one of these paralogs closely matches the sequence that one of the functional studies labeled as *HsOr259-L2* due to the fact that this sequence was not present in the old genome assembly but bore similarity to *HsOr259* (21). We have now named this gene *HsOr385*. Thus, we applied a combination of reciprocal BLAST searches and phylogenetic reconstruction (see below) to resolve the naming incongruencies. We have also changed paralog names of the HsOrXXX.X format to HsOrXXX, assigning previously unused gene numbers where necessary. Below is the summary of changes we introduced to HSAL70:

- *HsOr307*.*1* is renamed to *HsOr307*;
- *HsOr307*.*2* is renamed to *HsOr380*;
- *HsOr131*.*1* is renamed to *HsOr131*;
- *HsOr131*.*2* is renamed to *HsOr381*;
- *HsOr93*.*1* is renamed to *HsOr93*;
- *HsOr93*.*2* is renamed to *HsOr382*;
- *HsOr89*.*1* is renamed to *HsOr89*;
- *HsOr89*.*2* is renamed to *HsOr383*;
- *HsOr77*.*1* is renamed to *HsOr77*;
- *HsOr77*.*2* is renamed to *HsOr384*;
- *HsOr257*.*1* is renamed to *HsOr257*;
- *HsOr257*.*2* is renamed to *HsOr258*;
- *HsOr257*.*3* is renamed to *HsOr259*;
- *HsOr257*.*4* is renamed to *HsOr385*;
- *HsOr258* is renamed to *HsOr386*;
- *HsOr182*.*1* is renamed to *HsOr182*;
- *HsOr182*.*2* is renamed to *HsOr387*;
- *HsOr211*.*1* is renamed to *HsOr211*;
- *HsOr211*.*2* is renamed to *HsOr388*;
- *LOC105187615* is renamed to *HsOr303*;
- *HsOr378* and *HsOr379* names are swapped to remain consistent with Slone et al. (21).

We have named the new version of gene annotations HSAL71, and attached is the corresponding .gtf file as Supplementary Data 1.

Additionally, we have determined that some of the sequences used for functional characterization may be artifactual (20,21). For example, the gene characterized as HsOr263 appears to be a truncated version of HsOr348 rather than a unique gene (21). Sequence labeled as HsOr236 appears to be a hybrid between current HsOr236 and HsOr238 and was likely synthesized based on incorrect old genome assembly (21). We have added these details to the Figure 1A legend to denote such cases.

Nucleotide sequence of each *Or* was translated using TransDecoder v5.7.1 (Haas, BJ. https://github.com/TransDecoder/TransDecoder) by running TransDecoder.LongOrfs with default parameters, followed by TransDecoder.Predict with the --single_best_only flag to only keep one longest translation per gene. Resulting *Or* amino acid sequences were aligned using MAFFT v7.526 with default parameters (36). Next, the most informative sites were manually selected using Jalview v2.11.4.1 (37). Finally, a maximum likelihood tree was built using RAxML v8.2.12 with the following parameters: -f a -m PROTGAMMAAUTO -# 100 to perform 100 rapid bootstrap searches, 20 maximum likelihood searches, and return the best maximum likelihood tree (38). The obtained tree was rooted with *Orco* as an outgroup using FigTree v1.4.4 (https://github.com/rambaut/figtree) and plotted using the ggtree v3.10.0 library in R (39). Wherever possible, tree branches were reordered to match the alphabetical order of subfamilies by applying rotateConstr() function from the package ape v5.8 (40).

### Molecular Biology and Transgenic *Drosophila*

The 23 HsOr genes used in this study were obtained using one of the following two approaches. One approach began with extraction of RNA from resected and manually disrupted antennal tissue from *H. saltator* using TRIzol reagent (Invitrogen, ThermoFisher Scientific, Waltham, MA, USA). Antennal cDNA was then synthesized using Superscript II Reverse Transcriptase and the oligo(dT) primer (Invitrogen). PCR primers for each *HsOr* cloned with this method are listed in Supplementary Table S1. The forward primers were synthesized with additional 5’ CACC and amplified the predicted full-length HsOr gene sequences from antennal cDNA. Amplified *HsOr* genes were gel-purified and cloned into the entry vector via the pENTR™/D-TOPO™ Cloning Kit (Invitrogen, ThermoFisher Scientific, Waltham, MA, USA). Alternatively, our second approach synthesized HsOr genes with flanking attL recombination sites that were ligated into the pUC57-Kan vector (Genscript, Piscataway, NJ, USA) to generate entry vector. *HsOr* gene sequences were predicted full-length sequences supported by transcriptomic data (16). *HsOrs* in entry vectors from both approaches were transformed, purified, and sequenced, and then subsequently subcloned into pUASg.attB (gift from Konrad Basler, University of Zurich) with LR Clonase II enzyme (Invitrogen, ThermoFisher Scientific, Waltham, MA, USA). Resulting pUASg.attB-HsOr plasmids were then injected into *Drosophila melanogaster* embryos from the attp40w line with a genetically encoded phiC31 integrase (Rainbow Transgenic Flies Inc., Camarillo, CA, USA). Injected flies were then crossed with a CyO and TM3 balancer line, analyzed by PCR to confirm HsOr integration, and then crossed with the Orco-GAL4 driver line (Bloomington *Drosophila* Stock Center #23292) to generate homozygous individuals for electrophysiological screening.

### Electrophysiology

Single-sensillum recordings were performed using flies that were 4-7 d post-eclosion. Experimental fly genotypes were *w*; *w*^*+*^, *UAS-HsOr; w*^*+*^, *Orco-GAL4* and control flies were *w*; *+; w*^*+*^, *Orco-GAL4*. Within the constraints of fly line availability, trials were randomized. Extracellular recordings from ab2 sensilla were performed on mounted flies as described previously (20). For each recording, the target ab2 sensillum type was confirmed using a 6-odor diagnostic panel. The diagnostic odors were diluted 100-fold in paraffin oil and consisted of paraffin oil, 2-heptanone, ethyl acetate, geranyl acetate, (*E*)-2-hexenal, and racemic 1-octen-3-ol. 20 μL of each odor solution was loaded onto cotton-plugged Pasteur pipette odorant cartridges. Following diagnostic screening, a panel of 40 HCs, including a solvent control (pentane), were diluted to 10 mM in pentane and 20 nmol (2 μL) were applied to the interior of unplugged glass Pasteur pipettes (∼3 cm from open end of the pipette) on a location marked on the outside of the pipet. The panel included several unsaturated and branched HCs that were synthesized as previously described (20,41–43). The entire screening panel is listed in Supplementary Table S2.

Each delivery cartridge with HC was heated for 1 s using a handheld butane torch with the blue tip of the flame centered on the marked location where HC was deposited. Using a CS-55 Stimulus Controller (Syntech, Buchenbach, Germany), airflow through a blank cartridge was switched to the test odorant cartridge for 1 s (6 mL/s) into the continuous 20 mL/s humidified airstream. There was a 200 ms delay between the initiation of the odorant puff and the odorant reaching the antenna due to the length of the air delivery tube. Therefore, spikes were counted manually for both the 1 s window before stimulus application and a 200 ms window between 0.2 and 0.4 s of the 1 s stimulus. The Δspikes/s was calculated by subtracting pre-stimulus spike frequency and then normalized to the response of the heated solvent (pentane). Subsequent dose-response assays were performed on the HsOr-ligand pairs which showed responses above the 30 spikes per second threshold.

### Data Visualization of 70 HsOr Response Profiles

Figures 5 and 6 were generated using R (v4.5.0; R Core Team 2025) combining all HsOr functional data. Any missing response values due to absence from the HC screening panel or lack of expression data were eliminated from the data set. For Figure 5, response values were grouped by subfamily and summed. Each combined response value was visualized in Figure 6A and the average HsOr response across all HsOrs tested in Figure 6B. In Figure 6C, each combined response value was multiplied by the fragments per kilobase of transcript per million mapped reads (FPKM) expression value from a worker ant for a given HsOr. In Figure 6D, the average receptivity value was calculated from the weighted response values in Figure 6C. A Shiny App for the HsOr Response Database (https://pask-lab.shinyapps.io/HsOr_Response_Database/) was made by joining the combined response values with SEM values from the raw data set (44). Three tabs were made to visualize response values by receptor, hydrocarbon, and subfamily separately. The response curves, dose-response curves, and heat maps were generated in GraphPad Prism 10 (GraphPad Software).

## ACKNOWLEDGMENTS

We thank the Bucknell University and Middlebury College vivarium staff for assistance in maintaining ant colonies and fly stocks. We thank Laurence J. Zwiebel, Anandasankar Ray, and Jesse D. Slone for sharing previous HsOr datasets for our extended analyses. We also thank Eamon McMahon for fabrication of our Faraday cages and the Middlebury Makerspace for 3D-printing and laser cutting of electrophysiology equipment.

