## Supplementary Table S1 for "Sensitivity to Cuticular Hydrocarbons Across the Odorant Receptor Family in the Indian Jumping Ant"

| Target Gene | Forward Primer (5' to 3') | Reverse Primer (5' to 3') |
|---|---|---|
| HsOr219 | CACCATGAGTCTATCAATTGTTTATATCTTGGGC | CTAAGCGGCTGAGCTTTGAAGCACC |
| HsOr228 | CACCATGGAACTGCTTTCGTTGAATTTTTTCA | TCAGGACTGTTGCAGCACGTTGA |
| HsOr240 | CACCATGCACATACTTTCATTGACTTTTGCTCT | CTAAGAATCTTTCAGAAAGTTGTACGTCGAATA |
| HsOr191 | CACCATGTGTTTCAGCCAAGTTGTCTTC | TTATTTAGACTCGTCCAGTGATTGCAAA |
| HsOr213 | CACCATGGCGAGAAAAGTAACTCCAAAAGT | TTAACTTTTAGTTGCGTCGTCCTC |
| HsOr42 | CACCATGTCGTCGGTCGACCGT | TTACGCCGCTACCATTCGAATGAAA |
| HsOr16 | CACCATGCAACAGAGCATCCAGCTGAA | TTACGAAGTTAATGTGCGTAACATGTTC |
| HsOr62 | CACCATGGACTCGAACGCGCAGTG | TCACGTTTCTAACATCACTCTTAATACCGAC |
| HsOr72 | CACCATGCCGGATGATCGCTG | TTATTCGTTCATCAACGTTACTCGCAAG |
| HsOr70 | CACCATGACAAGTGAACGATGGAACGA | TTATGTTTCTACCATTACTCGAAGTACTGACAA |
| HsOr187 | CACCATGAAAGTGAACCGGGTGG | TCACATAAAGTTGCGCAATATGGATAAGTA |
| HsOr186 | CACCATGGCATATGCGAATTTTTACGAAGTC | TCACAACAGAGTACGCAGCACC |
| HsOr180 | CACCATGTCGACGTTGGGGCTC | TTACAAAAAAGAACGCAGCACCGACA |
| HsOr115 | CACCATGCTAAAAATCATTGGTTTGTGG | TTATGAATTTTGCTTGGCAAGTAACA |
| HsOr129 | CACCATGGACTGTCTGAGCACCTTTG | TCATTGCATCGCCATTAATACCGAGATG |
| HsOr139 | CACCATGGCGTTGAGTACAGCTCAGG | TCAATACATCGCCAAGAGCACCGA |
| HsOr152 | CACCATGTCGTTCATCCTGATCGTG | TCAATACGTCGCATTCAAGACGGAC |

**Supplementary Table S1. Primer Sequences used to clone full-length HsOrs.** All genes above were cloned via the pENTR™/D-TOPO™ Cloning Kit (Invitrogen, ThermoFisher Scientific, Waltham, MA, USA). All forward primers include a 5' CACC to facilitate directional cloning into the entry vector.
