## Supplementary Table S2 for "Sensitivity to Cuticular Hydrocarbons Across the Odorant Receptor Family in the Indian Jumping Ant"

| Order in panel | Compound Name | Common Name | Supplier |
|---|---|---|---|
| **1** | Paraffin Oil | | Avantor |
| **2** | 2-Heptanone | | Alfa Aesar |
| **3** | Ethyl Acetate | | Acros Organics |
| **4** | Geranyl Acetate | | Alfa Aesar |
| **5** | (*E*)-2-Hexanal | | Alfa Aesar |
| **6** | 1-Octen-3-ol | | Alfa Aesar |
| **7** | Pentane | C5 | Fisher Scientific |
| **8** | Decane | C10 | Alfa Aesar |
| **9** | Undecane | C11 | Acros Organics |
| **10** | Dodecane | C12 | Alfa Aesar |
| **11** | Tridecane | C13 | Acros Organics |
| **12** | Tetradecane | C14 | Acros Organics |
| **13** | Pentadecane | C15 | Acros Organics |
| **14** | Hexadecane | C16 | Alfa Aesar |
| **15** | Heptadecane | C17 | Acros Organics |
| **16** | Octadecane | C18 | Acros Organics |
| **17** | Nonadecane | C19 | Alfa Aesar |
| **18** | Eicosane | C20 | Alfa Aesar |
| **19** | Heneicosane | C21 | Acros Organics |
| **20** | Docosane | C22 | Acros Organics |
| **21** | Tricosane | C23 | Alfa Aesar |
| **22** | Tetracosane | C24 | Alfa Aesar |
| **23** | Pentacosane | C25 | Sigma-Aldrich |
| **24** | Hexacosane | C26 | Alfa Aesar |
| **25** | Heptacosane | C27 | Alfa Aesar |
| **26** | Octacosane | C28 | BeanTown Chemical |
| **27** | Nonacosane | C29 | Sigma-Aldrich |
| **28** | Triacontane | C30 | Acros Organics |
| **29** | Hentriacontane | C31 | TCI |
| **30** | Dotriacontane | C32 | Acros Organics |
| **31** | Tritriacontane | C33 | TCI |
| **32** | Tetratriacontane | C34 | Acros Organics |
| **33** | Pentatriacontane | C35 | TCI |
| **34** | Hexatriacontane | C36 | Alfa Aesar, TCI |
| **35** | Heptatriacontane | C37 | Sigma-Aldrich |
| **36** | 2-Methyloctacosane | 2-MeC28 | J.G. Millar |
| **37** | (Z)-9-Nonacosene | 9-C29:1 | J.G. Millar |
| **38** | 3-Methylnonacosane | 3-MeC29 | J.G. Millar |
| **39** | 13-Methylnonacosane | 13-MeC29 | J.G. Millar |
| **40** | 15-Methylnonacosane | 15-MeC29 | J.G. Millar |
| **41** | 2-Methyltriacontane | 2-MeC30 | J.G. Millar |
| **42** | (Z)-9-Hentriacontene | 9-C31:1 | J.G. Millar |
| **43** | 5-Methylhentriacontane | 5-MeC31 | J.G. Millar |
| **44** | 13-Methylhentriacontane | 13-MeC31 | J.G. Millar |
| **45** | 15-Methylhentriacontane | 15-MeC31 | J.G. Millar |
| **46** | 13,23-Dimethylheptatriacontane | 13,23-DiMeC37 | J.G. Millar |

**Supplementary Table S2. Hydrocarbon panel used for functional characterization of HsOrs.** The first 6 odorants are the diagnostic panel used to confirm ab2 sensillum identity. Pentane (7) is the solvent for the other hydrocarbons and it is the vehicle control. The test panel compounds used for analysis of hydrocarbon response are compounds 8-46.
